# Socio-demographic correlates and clustering of non-communicable diseases risk factors among reproductive aged women of Nepal: Results from Nepal Demographic Health Survey 2016

**DOI:** 10.1101/669556

**Authors:** Bihungum Bista, Raja Ram Dhungana, Binaya Chalise, Achyut Raj Pandey

## Abstract

Globally, Non Communicable Diseases (NCDs) are the major killer diseases, majority of which are attributable to common risk factors like smoking, alcohol intake, physical inactivity and low fruits/vegetable consumption. Clustering of these risk factors increases the risk of developing NCDs. The occurrence of NCDs among women is alarmingly high, and this invites impact on upcoming generation too. So, this study aimed to assess the prevalence and clustering of selected risk factors and their socio-demographic determinants in Nepalese women using Nepal Demographic and Health Survey (NDHS) 2016 data.

NDHS applied stratified multi-stage cluster sampling to reach to the individual respondent for representing the whole nation. This study included analysis of data of 6,396 women of age 15 to 49 years. Chi-square test for bivariate analysis and multiple poisson regression to calculate adjusted prevalence ratio was applied.

A total of 8.91% participants were current smoker. Similarly, 22.19% and 11.45% of participants were overweight and hypertensive respectively. Around 6.02% of participants had a co-occurrence of two NCDs risk factors. Smoking, overweight and hypertension were significantly associated with age, education, province, wealth index and ethnicity. Risk factors were more likely to cluster in women aged 40-49 years (APR=2.95, CI: 2.58-3.38), widow/separated (APR=3.09, CI: 2.24-4.28) and Dalit) (APR=1.34, CI: 1.17-1.55).

This study found that NCD risk factors were disproportionately distributed by age, education, socio-economic status and ethnicity and clustered in more vulnerable groups such as widow/separated, Dalit and Janajati.

## Introduction

Globally, non-communicable disease (NCDs) are the number one causes of death and disability. NCDs account for 41 million deaths each year and 85% of these deaths occur in low- and middle-income countries while nearly half of NCDs deaths (15 million out of 41 million) occur between the age of 30 and 69 years.[1] Cardiovascular diseases, cancers, diabetes, and respiratory diseases, also called the ‘Group of Four’ are responsible for 80% of all NCDs deaths. [1]

NCDs share the common risk factors such as low intake of fruit and vegetables, low level of physical activity, tobacco use, harmful use of alcohol, obesity, raised blood pressure, raised blood cholesterol and glucose. The co-occurrence of these risk factors in individual is known as clustering of risk factors. Clustering of risk factors is related with an increased risk of developing NCDs.[2,3] In context of Nepal, STEPS survey 2013, reported that 15.5% of general population and 11.4% of women had three or more risk factors of NCD in them.[4] Evidence show that women are more likely to experience the co-occurrence of behavioral and metabolic risk factors increasing the risk of NCDs among themselves and in future generation.[5-7] Similarly, compared to men, women experience fewer symptoms and show less apparent signs of certain NCDs like cardiovascular disease. They are thus less likely to be identified and treated or less likely to be the focus of disease prevention.[8]

NCDs have broader impact that varies from –impact on maternal to child health, individual to national level and physical burden to financial burden. Thus to tackle with NCDs, the best strategy is to identify and modify the behavioral risk factors that causes NCDs. This study, therefore, aims to assess the magnitude of selected risk factors, individually or in cluster and determines their socio-demographic distributions in Nepalese women.

## Methodology

This study is based on the data from the Nepal Demographic Health Survey (NDHS) 2016.NDHS is periodic survey that consist of a nationally representative sample. A detailed description of NDHS methodology is reported elsewhere.[9] Briefly, NDHS applied the stratified multi-stage cluster sampling to reach to the individual respondent. Firstly, 383 primary sampling units (PSU) (wards) were selected based on probability proportional to PSU size. Subsequently, 30 households per PSU (total 11040 households) were selected using an equal probability systematic selection criterion. The 2016 NDHS was first time included the measurements of biomarker information including blood pressure. Blood pressure and anthropometric measurements were only obtained from the systematically selected subsample of the total study participants. For this study, we have only included 6396 women aged between 15 and 49 years who had their blood pressure recorded.

### Data collection

#### Blood pressure

Trained enumerator measured blood pressure with UA-767F/FAC (A&D Medical, Tokyo, Japan) blood pressure machines. Enumerators took three readings of blood pressure at the interval of five minutes between each reading and averaged last two readings to get more accurate blood pressure level. Participants whose systolic blood pressure (SBP)at the level of 140 mmHg or higher and/or diastolic blood pressure (DBP) of ≥90 mm Hg or higher or currently taking antihypertensive medicines at the time of data collection were considered hypertensive.

#### Overweight

Weight in kilograms was divided by height in meters-squared to calculate BMI. Women having (BMI ≥ 25kg/m^2^) were categorized as ‘Overweight” and the remaining (BMI< 25kg/m^2^) were categorized as “Not overweight”.

#### Current tobacco use

Current tobacco use includes either daily or occasional smoking or use of smokeless tobacco (snuff by mouth, snuff by nose, chewing tobacco and betel quid with tobacco)

### Explanatory variables

For this study purpose, information related to socio-demographic variables including age of the participants, ethnicity, educational status, place of residence (rural/urban), province and ecological zone and wealth index were extracted from the NDHS original datasets.

### Statistical analysis

All analyses were performed on STATA 15.2 version using survey set command. All estimates were weighted by sample weights and presented with 95% Confidence Intervals. Prevalence estimates were calculated using Taylor series linearization. Chi-square test was used for bivariate analysis to test associations between covariates and dependent variables. Furthermore, multiple Poisson regression was used to calculate adjusted prevalence ratio (APR). The numbers of risk factors present within each participant (from 0 to 3) were counted to assess clustering of risk factors and analyzed using the Poisson regression.

### Ethical consideration

The 2016 NDHS ethical approval was sought from Ethical Review Board (ERB) of the Nepal Health Research Council (NHRC), Nepal and ICF Macro Institutional Review Board, Maryland, USA. Written informed consent was obtained from each participant before enrolling in the survey.

## Results

Just over half (53.95%) of the participants were of aged 15-29 years. Largest proportions (36.62%) of the participants were from Janjati group (indigenous group). One thirds (33.34%) had no formal schooling while 76.55% of the participants were married. Most of the participants belonged to Terai belt (49.89%) and rural areas (63.30%).Similarly, 22.43% and 20.92% of participants belonged to richer and richest wealth quintile. Most of participants were engaged in agriculture or were self employed.

## Distribution of non communicable diseases risk factors

### Current tobacco use

The prevalence of current tobacco use was 8.91%.Women aged 40-49 years (22.38%), no education (18.81%) and widowed/divorced/separated (29.07%) had the highest prevalence of current tobacco use among their respective categories “Table 2”. Similarly, current tobacco use was significantly associated with ecological zone, province, wealth index, occupation and ethnicity “Table 2”.

**Table 1:** Socio-demographic distribution of participants.

| Characteristics | un-weighted count | weighted percent |
| --- | --- | --- |
| **Age group** |  |  |
| 15-29 | 3,498 | 53.95 |
| 30-39 | 1,697 | 27.09 |
| 40-49 | 1,201 | 18.96 |
| **Educational status** |  |  |
| No education | 2,161 | 33.34 |
| Primary | 1,017 | 16.7 |
| Secondary | 2,324 | 35.48 |
| Higher | 894 | 14.48 |
| **Marital status** |  |  |
| Never in union | 1,305 | 20.73 |
| Married or living together | 4,919 | 76.55 |
| Widowed/divorced/separated | 172 | 2.72 |
| **Ecological region** |  |  |
| Mountain | 454 | 6.05 |
| Hill | 2,916 | 44.06 |
| Terai | 3,026 | 49.89 |
| **Residence** |  |  |
| Rural | 4,129 | 63.03 |
| Urban | 2,267 | 36.97 |
| **Province** |  |  |
| Province 1 | 909 | 16.84 |
| Province 2 | 1,051 | 19.94 |
| Province 3 | 853 | 22.07 |
| Gandaki | 803 | 9.81 |
| Province 5 | 988 | 16.87 |
| Karnali | 888 | 5.66 |
| Sudurpaschim | 904 | 8.82 |
| **Wealth index** |  |  |
| Poorest | 1,347 | 16.96 |
| Poorer | 1,304 | 19.11 |
| Middle | 1,319 | 20.57 |
| Richer | 1,319 | 22.43 |
| Richest | 1,107 | 20.92 |
| **Occupational status** |  |  |
| Did not work | 2,003 | 32.34 |
| Services | 863 | 15.01 |
| Agriculture/ self-employed | 3,196 | 46.88 |
| Manual | 331 | 5.77 |
| **Ethnic group** |  |  |
| Advantage group | 2,254 | 31.31 |
| Dalit | 851 | 12.56 |
| Janjati | 2,268 | 36.62 |
| Other | 1,023 | 19.51 |
| **Total** | **6,396** | **100** |

**Table 2:**
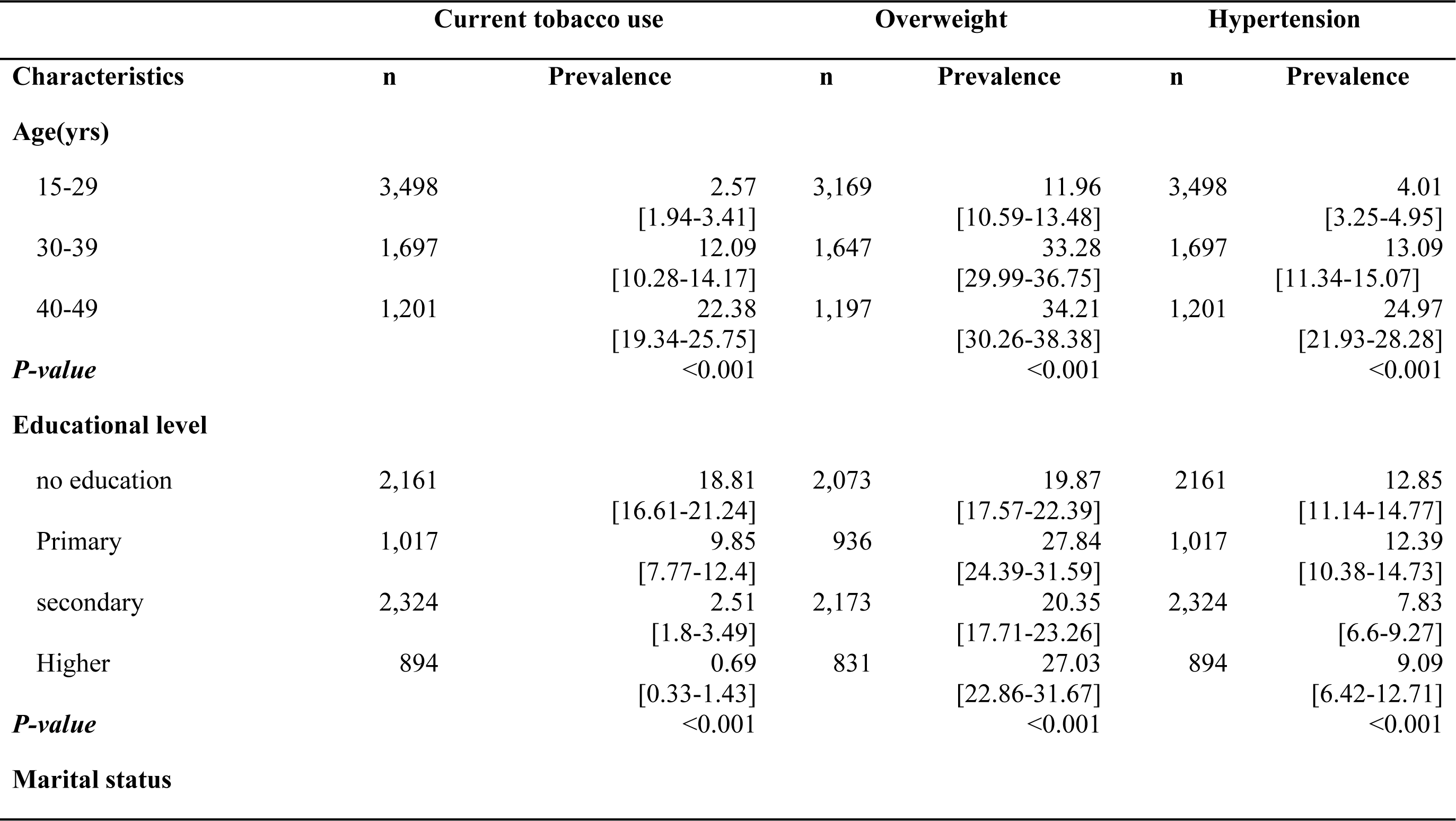
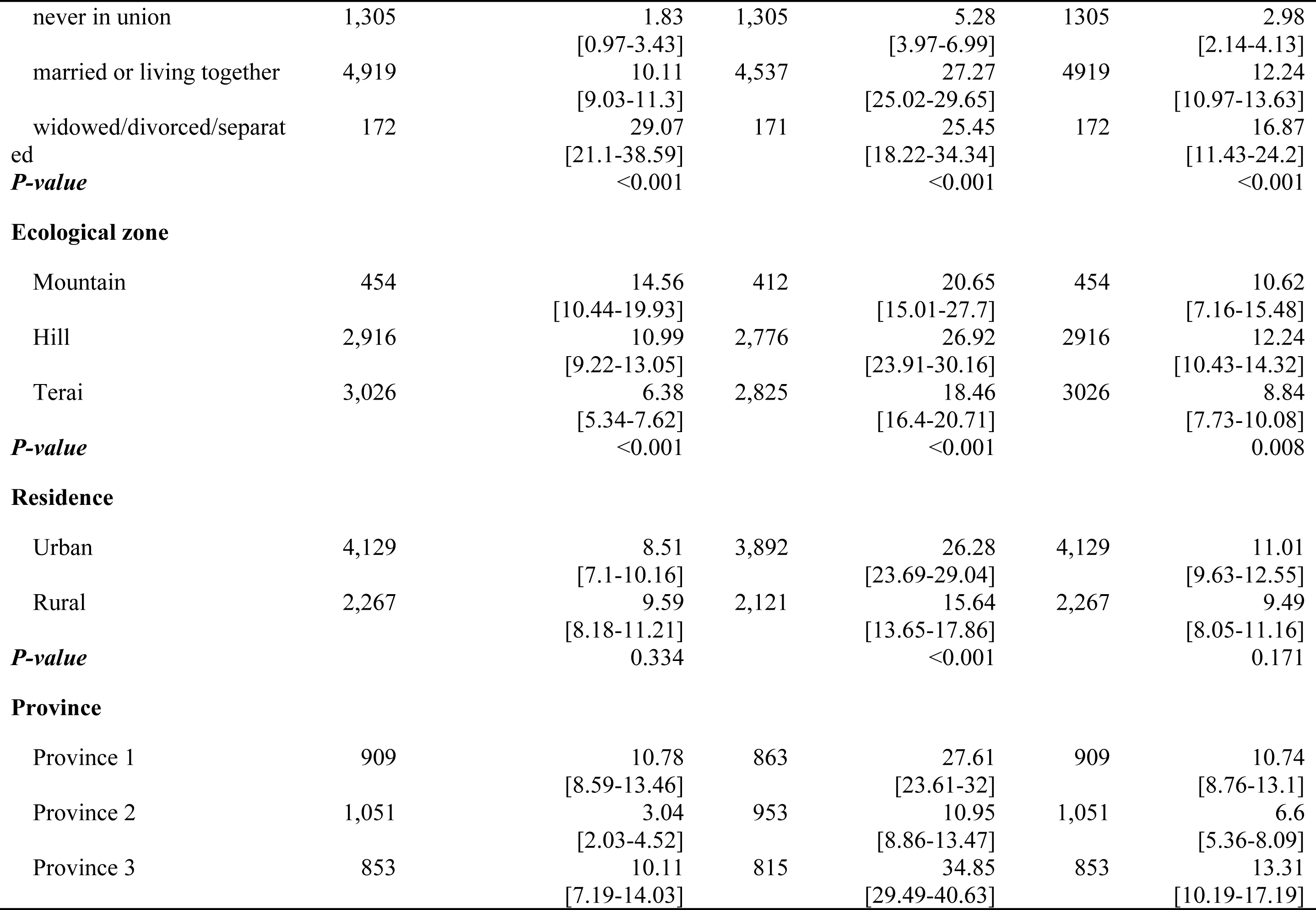
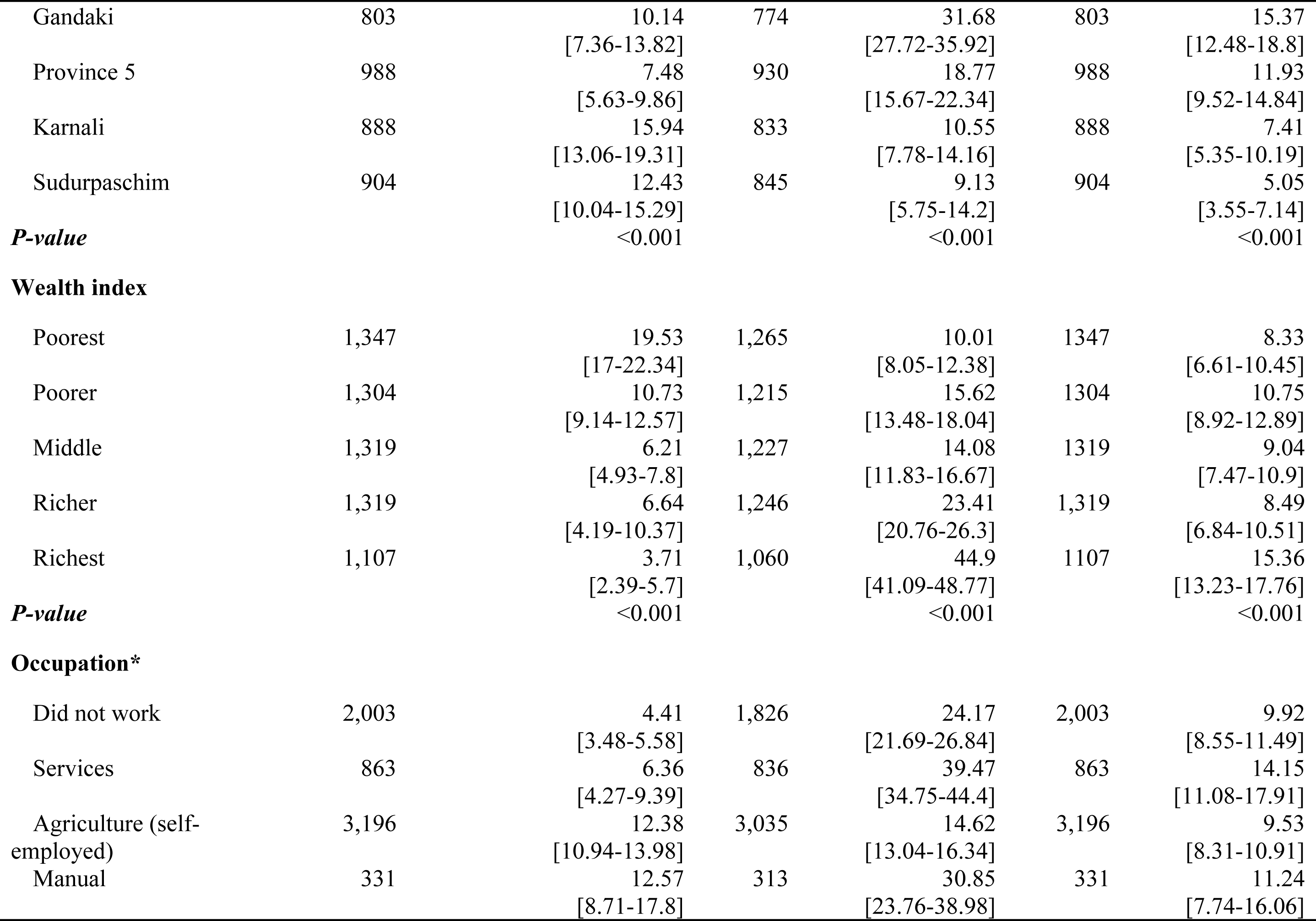
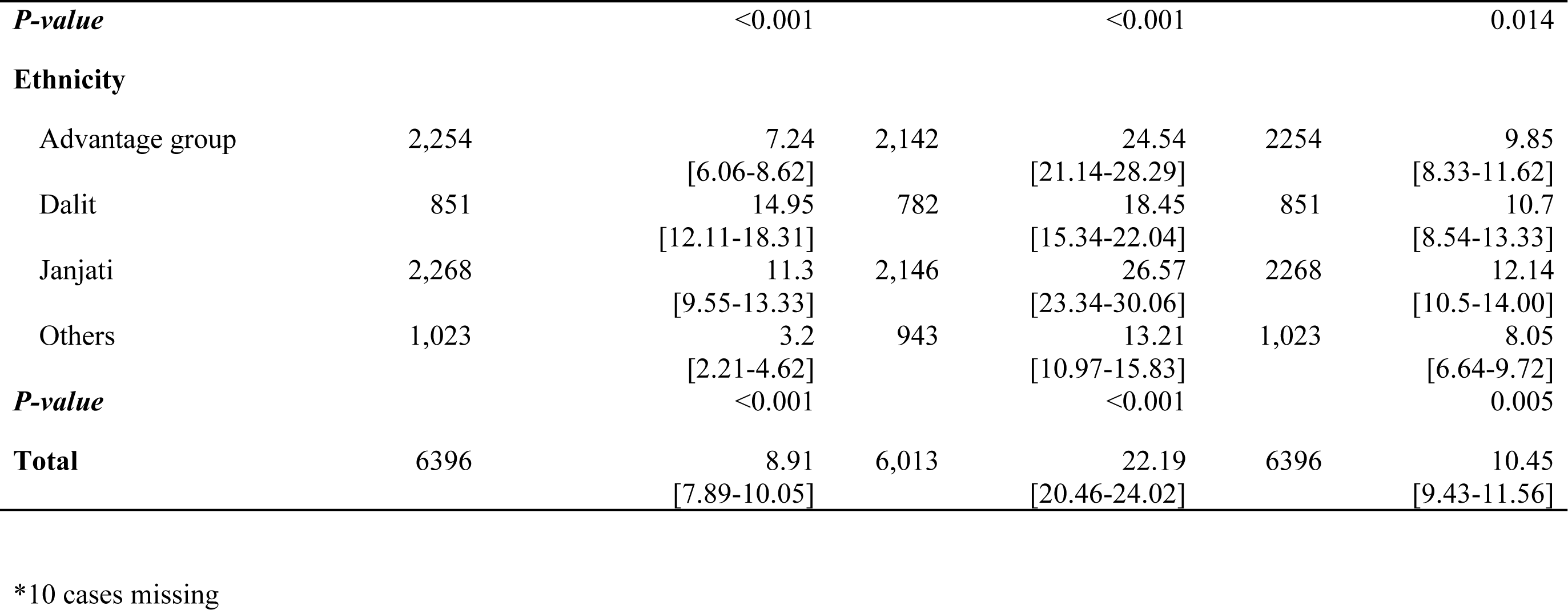
Prevalence (%) of non-communicable diseases risk factors among 15-49 years women.

**Fig 2:**
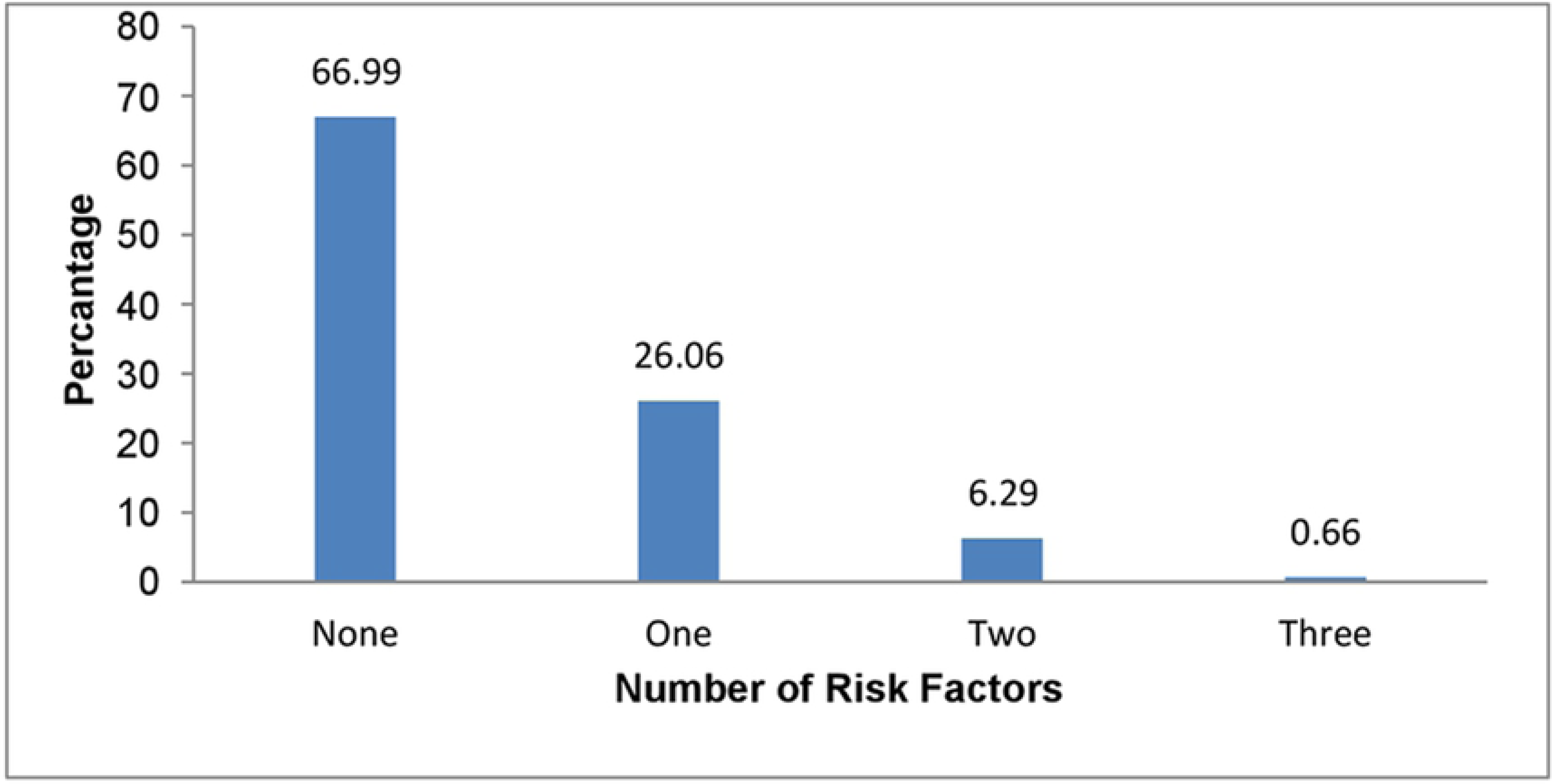
Prevalence of number of NCDs risk factors among participants. 26.08% of participants had one NCDs risk factors and 6.3% participants had two NCD risk factors (Fig 1)

### Overweight

The prevalence of overweight/obesity was 22.19%. The rate was significantly high in women aged 40-49 years compared to that of 15-29 (11.96%) years women “Table 2”. Similarly, prevalence of overweight significantly varied by education status “Table 2”. Compared to never union, prevalence of overweight is significantly high among married/ living together women (27.27%) or divorcee/widowed/separated (25.45%). Current tobacco use is also significantly associated with residence status, province, wealth index, occupation and ethnicity “Table 2”.

### Hypertension

Prevalence of hypertension was 10.45%. The prevalence of hypertension significantly varied by the age of the participants, where women aged 40-49 years had the highest rate of hypertension. Secondary education was significantly associated with higher prevalence of hypertension compared to primary and no education. Likewise, the rate of hypertension was also significantly different in province, wealth index, occupation and ethnicity “Table 2”.

## Multivariable analysis of socio-demographic characteristics with non-communicable diseases risk factors

### Current tobacco use

Women of age 30-39 years and 40-49 years were 2.56 and 3.70 times more likely to be tobacco user than that of 15-29 years old women “Table 2”. Similarly, educated women were less likely to be tobacco user (APR primary: 0.71, APR secondary: 0.28, APR Higher: 0.09) than that of uneducated women. Widowed/divorced/separated women were 2.03 times more likely to be tobacco user than that of women who were never in union. Furthermore, women residing on province 2 (APR: 0.28) and province 3 (APR: 0.64) were less likely to be tobacco user in comparison to province 1 women. Similarly, poor women (APR: 0.69) were more like to be smoker than that of poorest women. Dalit women were 1.68 times more likely to be tobacco user in comparison to advantage women.

### Overweight

Women of age 30-39 years were 1.85 and 1.97 more likely to overweight in reference to 15-29 years “Table 2”. Similarly, married and single women were 4.02 and 3.29 respectively times more likely to be overweight than that of never in union women. Further, more women residing in province 2, province 5, Karnali and Sudurpaschim were less likely to overweight in comparison to province 1.Regarding wealth, as the gradient of wealth increases women were more likely to be overweight in comparison to poor women. Women involved self-employed agriculture were less likely to be overweight in reference to who didn’t have work.

### Hypertension

Women of 40-49 years and 30-39 years were 1.97 and 1.85 times more likely to be hypertensive in comparison to 15-29 years women “Table 2”. Primary educated women were 1.27 times more likely to be hypertensive in comparison to uneducated women. Married and single (widowed/separated/divorced women) were 4.02 and 3.29, respectively, times more likely to be hypertensive than that of never in union women. Similarly women residing in province 2 province 4 province 5 province 6 and province 7 were less likely to hypertensive in comparison to province 1 women. Richest women were 3.38 times more likely to be hypertensive than that of poorest women. Women whose occupation was agriculture were less likely to be hypertensive in comparison to who didn’t have work.

### Clusteringof NCDs risk factors

Women of 40-49 years age group were 2.95 more times likely to have NCD risk factors than that of 15-29 years of women “Table 3”. Women who had pursued secondary level of education were 0.87 times less likely to have NCD risk factors. Married and widowed/divorced/separated women were 2.91 and 3.09 times more likely to have NCD risk factors. Similarly, richest women were 1.5 times more likely to suffer from NCDs risk factors in comparison to poorest women. Furthermore, women employed in agriculture sector were 0.83 times less likely to suffer from NCD risk factors than women who were not employed. Regarding ethnicity, Dalit women were more likely to have NCD risk factors in comparison to advantage group.

**Table 3:**
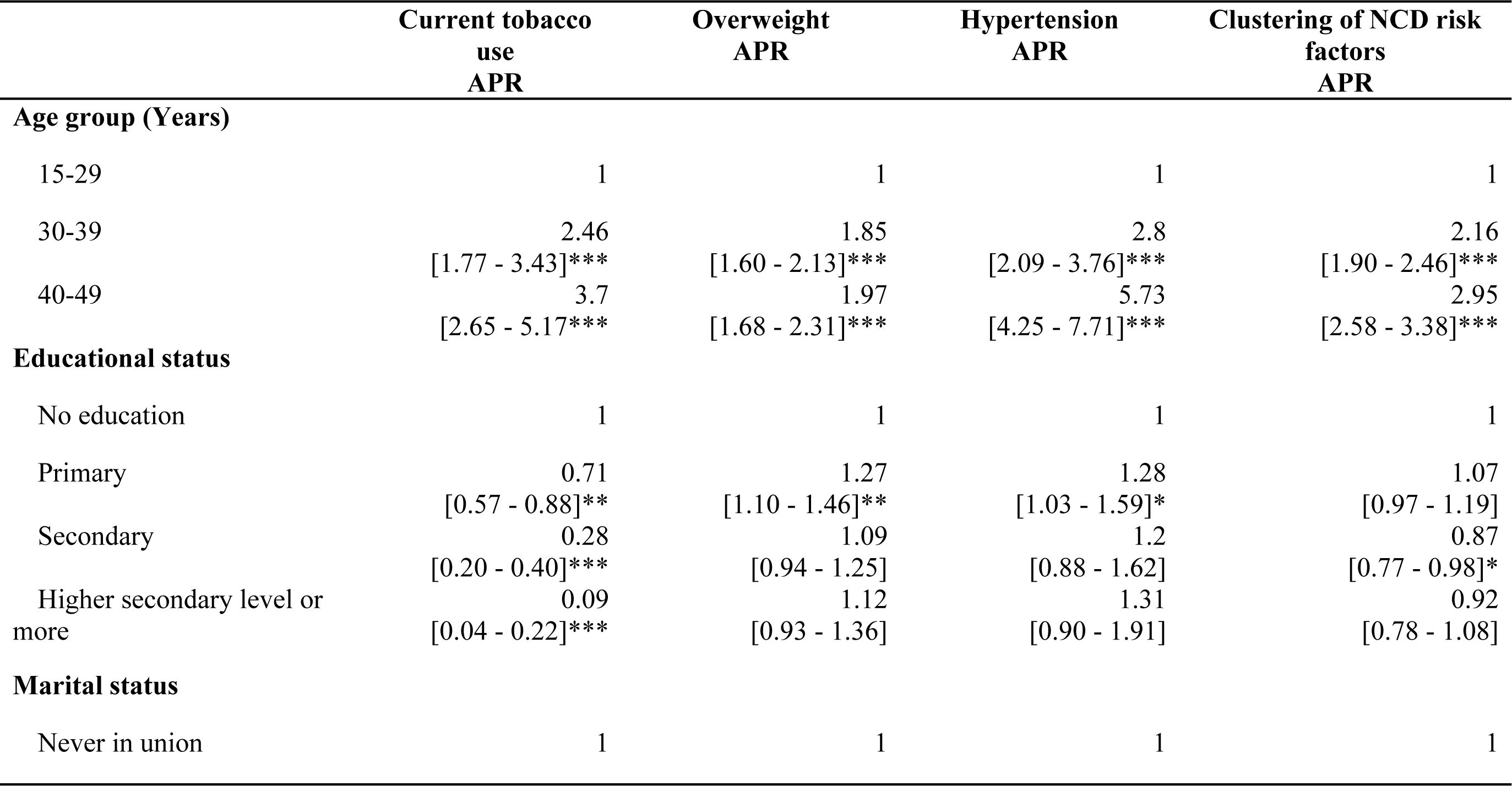
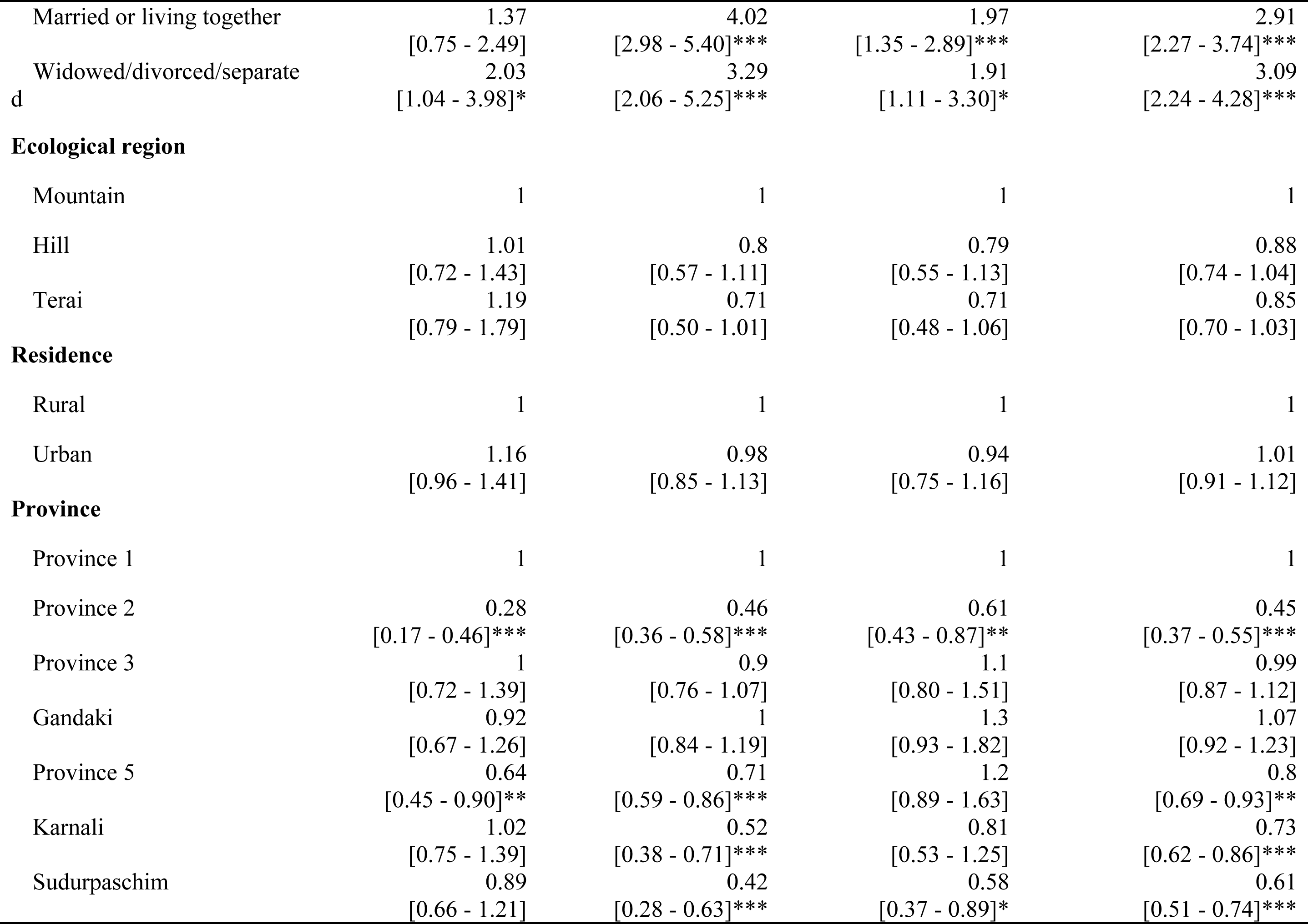
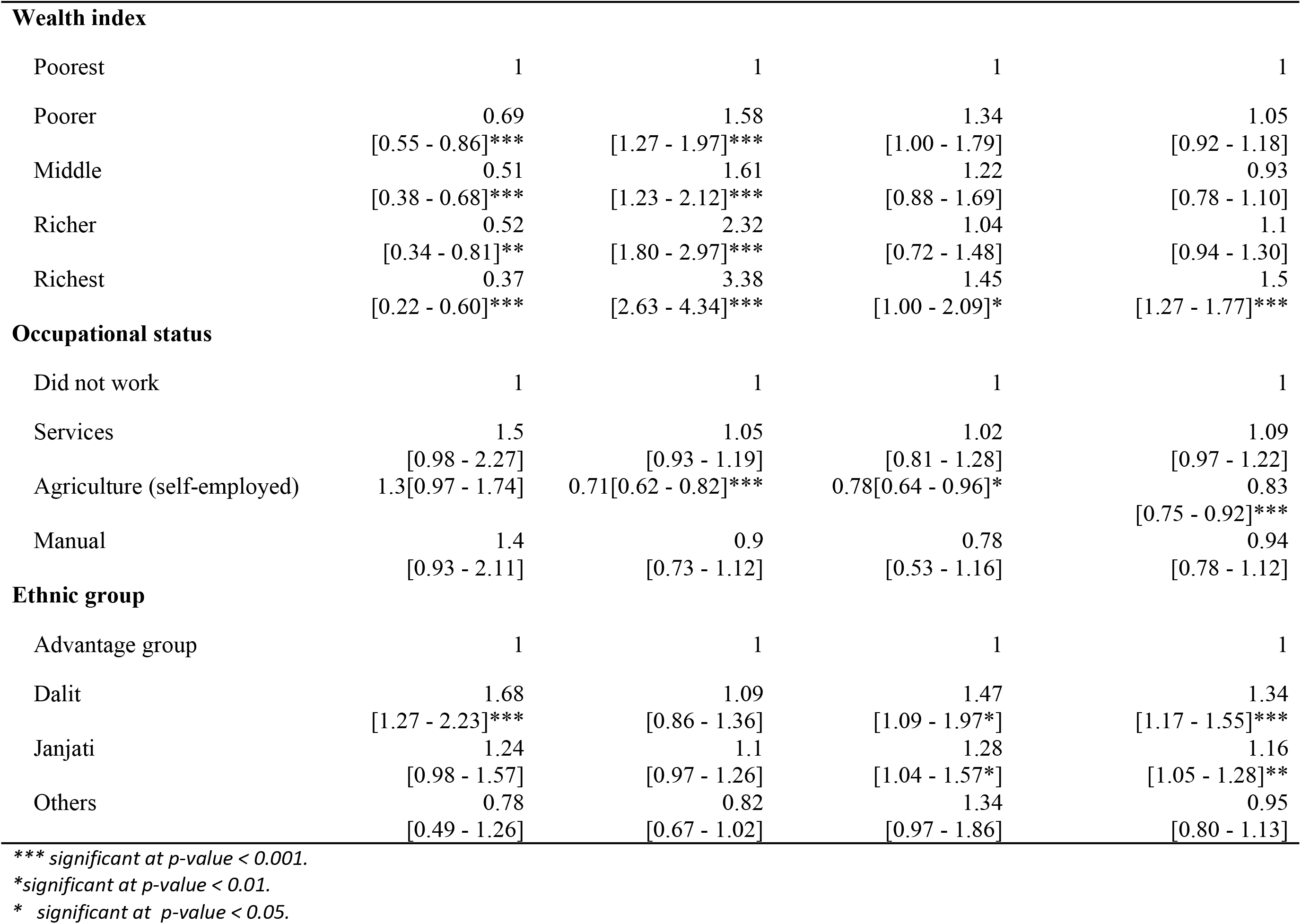
Relationship of socio-demographic characteristics with non-communicable diseases risk factors.

## Discussion

NCDs have different consequences for women in comparison to men.[10] In resource challenged setting like Nepal, diagnosis and care for NCDs are less accessible and affordable to women due to limited health infrastructure and human-resource capacity. As a result, NCDs are often detected at the late stage that invites women for a premature death. So, this study aimed to identify at risk women to possess NCDs risk factors. This information could be useful in designing preventative strategies against NCDs risk factors.

### Tobacco use

Our study demonstrated that the proportion of tobacco use was nearly 3 fold higher in 30-40 years age group women. This finding is in line with previous studies conducted across different countries.[11,12] Women aged between 30 and 40 years are likely to possess the adolescent children thus maternal smoking could significantly contribute to tobacco use in young adolescent.[13] High prevalence of tobacco use in the women with childbearing age is also critical in terms of adverse maternal and child health outcomes in perinatal period.[14]

Our study showed a negative association between smoking and education; the prevalence ratios among the participants having secondary and higher education being lower than those having no education. This finding is similar to that of previous studies.[11,12]

In this study, tobacco use was higher in divorced women than married women. Similar kind of evidence was reported in other studies, especially related with tobacco smoke.[11,15], which suggest that the death of loved ones can encourage women to opt smoking with intention of coping stress arising from the death of intimate partner.[16,17]

Unlike the findings from other national studies, [11,18] it is interesting to note that prevalence of smoking did not vary significantly between rural and urban participants. However, it should be taken into consideration that large number of geographical cluster previously considered as rural areas have recently been upgraded as urban that somehow makes the comparison difficult. Generally, people with low socioeconomic status are likely to use tobacco, probably due to lack of proper social support environment against tobacco use.[19] Our study also demonstrated the same finding that the poorest has the highest proportion of tobacco user across all hierarchies of wealth quintile. Evidence suggest that increase in taxation can be other effective strategy in controlling tobacco as there seem to be high price elasticity particularly low and middle income countries like Nepal.[20] Around 10% increase in price is found to reduce smoking by about 8% in low- and middle-income countries and by 4% in high-income countries.[21]

This is the first study to repute and compare smoking prevalence by Provinces of Nepale. Findings show that women from province 2 and province 5 were less likely to use any form of tobacco in comparision to women residing in other provinces. Province 2 and a major portion of province 5 share a similar kind geographical terrain i.e plain, where the media accessibility among women is high in comparison to other.[22]

The observed discrepancy in smoking prevalence by socioeconomic status may be related to less successful quiet attempts in disadvantaged groups.[19] Population level interventions such as smoke free legislation and mass media campaign tailored to the need of disadvantaged communities are thus need to reduce high smoking prevalence in the disadvantaged groups.

### Obesity

The current study found that the likelihood of being overweight/obesity is influenced by age, married marital status and wealth status. There is an increase in prevalence of overweight with increase age, this finding is line with findings of STEPS Nepal 2013 survey and studies from Bangladesh[2,4]. Similarly, this study found that married women were more likely to be overweight in comparison to unmarried women. Broadly, literatures explain that smoking and habit of looking attractive is related with obesity. [23,24] However, our another findings related with widowed/divorce were different than established crisis model. This model explains that stresses linked to marital disruption can invite psychological, physiological, and social consequences that might lead to weight loss.[24,25] However, this model explains that weight loss is short lived, individuals are expected to gain weight to their new social and economic environment.

Furthermore; wealthy women are on risk of getting overweight in comparison to poor women. The increased risk of getting overweight among wealthy and elderly women may be due to reduced level of physical activity with increased age and wealth status. Maternal obesity is a public health concern. The prevalence of overweight/obesity in reproductive age women is nearly tripled from 9% in last ten years in Nepal.[9,22,26].With the global rise in maternal obesity,[27] more mother and child are at risk of dying. There is a growing evidence that maternal obesity can substantially interfere in fetal development and determines the long term health of the offspring.[28] Similarly, it is also a major risk factor for gestational diabetes, preeclampsia and pregnancy induced hypertension in women. [29,30]

On the other hand, societal and nutritional changes due to economic growth and globalization of food market might have contributed the rising obesity rates. The lower level of education and health literacy among poor also contributes to difficulty in purchasing less energy dense food such as fruits and vegetable.

Further research in the area of food security, dietary pattern and physical activities by socio economic status is needed to rule out causes of obesity for women from low socioeconomic background versus high socioeconomic status. Intervention to tackle obesity requires targeting social and economic factors.

### Hypertension

The prevalence of hypertension seems to increase with increasing age, which is in line with results of secondary data analysis of Nepal STEPS survey 2013 and other evidences as well.[31-33] However, age being a non-modifiable risk factor, hypertension control initiatives should focus on lowering other modifiable risk factors that can be useful in countering the effect of increasing age.

The prevalence of hypertension was significantly higher among the richest segment of study participants. Similarly, chance of getting hypertension is significant in richest segment of population. Similar type of evidence was observed i.e higher prevalence of hypertension among richest segment in general population of Bangladesh.[33] It could be because of lower level of physical activity associated with involvement in more sedentary type of occupation, consumption of red meat, smoking and alcohol consumption.

Compared to province 1, province 7 have lower prevalence of hypertension. It can be due to differences in level of physical activity associated with occupational practices, dietary pattern, and differences in established risk factors of hypertensions like smoking and alcohol consumption.

### Clustering of NCDs risk factors and its implication

Our study suggested that clustering of NCDs risk factors increases with growing age, among well-off, and in Dalits and Janajatis - known as the disadvantaged ethnic groups in Nepal. Previous studies have also revealed that clustering of risk factors becomes increasingly common with increasing age [2,3,34]. A multi country study from Bangladesh, Vietnam, India, Indonesia and Thailand shows that there is increase in clustering of risk factors with increasing age among females.[35] As Nepal has been witnessing rapid increase in life expectancy and median age of the population, the problems can escalate in coming years.[36] Country may need additional investment in prevention as well as long term care for NCDs to cater the need of geriatric population. Moreover, NCDs are considered to have serious impact in economic growth of the country reducing it by almost 5–10%.[37]

Similarly, this study depicts the odds of clustering of NCDs risk factors higher among wealthiest women. This findings is similar to that of secondary data analysis of national STEPS survey from Bhutan.[38] Clustering of more NCDs risk factors in wealthy group can be linked with adoption of sedentary lifestyle among wealthy women. Furthermore, from provincial point of view provincial 3 and Gandaki province is not related with clustering of NCDs risk factors, however, other provinces had reduced odds of clustering of NCDs risk factors. This can be again viewed from the perspective of sedentary lifestyle with reference to urbanization. In comparison to province 1, province 3 and Gandaki province, other provinces are less urbanized that reduces odds of adoption of sedentary lifestyle. Ultimately, this might have contributed in reducing odds of clustering of NCD risk factors among women residing in province 5, Karnali and Sudurpaschim.

In contradiction to the study in Bangladesh, which revealed an increase in clustering of risk factors with increasing educational level, however our study shows that women who have secondary level of education had lower risk of clustering of NCDs risk factors. [35] The difference in evidence may be due to difference in NCDs prevention and control contents in secondary level education.

Furthermore, women involved in agriculture (self-employed) sector have low odds of clustering of NCDs risk factors. Generally, self-employed agriculture work is expected to increase the vigorous physical activity. Vigorous physical activity is a protective factor against obesity and it is expected to lower down the risk of clustering NCDs risk factors.[39]

As the burden of NCDs is increasing, evidence on clustering of NCDs risk factors is useful form the perspective of allocating and mobilizing resources in public health programme. The clustering of NCDs risk factors in a particular group indicte higher chances of NCDs burden on that particular group. This situation creates public health challenge; however, it can be also be an opportunity to tailor intervention for specific group of population to prevent the burden of NCDs. The limitation of this study is the nature of study i.e cross-sectional design that limit to establish causality. Similarly, all NCD risk factors like physical inactivity, fruit and vegetable intake, cholesterol level related information was not taken main survey. That had limited us to understand completed picture of NCDs risk factors among women.

## Conclusion

Overweight is the common NCD related risk factors among the 15-49 years women. The occurrence of NCDs related risk factors is higher in higher age group. However, in case of relationship of smoking with respect to wealth quintile relationship is inverse. Similarly, study reveals that chances of clustering of NCDs related risk factors get increases with increasing age. Furthermore, chances of clustering of NCDs risk factors are higher on disadvantaged ethnic group and richest women.

